# Engineering a model cell for rational tuning of GPCR signaling

**DOI:** 10.1101/390559

**Authors:** William M. Shaw, Hitoshi Yamauchi, Jack Mead, Glen-Oliver F. Gowers, David Öling, Niklas Larsson, Mark Wigglesworth, Graham Ladds, Tom Ellis

## Abstract

G protein-coupled receptor (GPCR) signaling is the primary method eukaryotes use to respond to specific cues in their environment. However, the relationship between stimulus and response for each GPCR is difficult to predict due to diversity in natural signal transduction architecture and expression. Using genome engineering in yeast, we here constructed an insulated, modular GPCR signal transduction system to study how the response to stimuli can be predictably tuned using synthetic tools. We delineated the contributions of a minimal set of key components via computational and experimental refactoring, identifying simple design principles for rationally tuning the dose-response. Using four different receptors, we demonstrate how this enables cells and consortia to be engineered to respond to desired concentrations of peptides, metabolites and hormones relevant to human health. This work enables rational tuning of cell sensing, while providing a framework to guide reprogramming of GPCR-based signaling in more complex systems.

## Introduction

G protein-coupled receptors (GPCRs) are widely represented in most lifeforms and comprise the largest family of signaling proteins in humans, with over 800 members detecting structurally diverse agonists^1,2^. Their abundance and ubiquity to all cell types makes them one of the most important signaling pathway classes in healthcare, but also one of the most complex^3,4^. Multiple types of G-protein based signaling are seen and the downstream signal transduction to activate gene expression is typically complex and intertwined with other pathways^5,6^. The nature of signal transduction through the pathway also depends on many different factors, including the stoichiometry of the signaling proteins, the presence of inherent feedback mechanisms, and even cellular history^7^. Altogether, this makes it difficult to delineate receptor and signaling properties simply from measuring the activation of downstream targets^8^. It also makes it a major challenge to predict how changes in the levels of pathway components, for example due to different environments or mutations, can affect the performance of a given signaling pathway.

The most studied example of a eukaryotic GPCR signaling pathway is the pheromone response pathway of *Saccharomyces cerevisiae*^9^, having been the focus of significant efforts from systems biology to model its actions via quantification of its behavior^10,11^. To understand this pathway, researchers have parsed the contributions of numerous studies that have perturbed the dose-response and dynamics of the native system by changing growth conditions, by protein mutagenesis, or via traditional gene overexpression or knockout methods^12,13^. While these efforts have helped to build our best picture of the events required for the transduction of signal from agonist to gene activation, inability to control the whole pathway in these experiments has meant that a complete system for exploring the dose-response relationship has not yet been achieved^13^.

*In silico* approaches typically model a system by concentrating only on the key components and varying important parameters of these such as their expression levels, while removing other non-key interactions from consideration^14,15^. With advanced genome engineering and synthetic biology tools available, it now becomes possible to take an equivalent modelling approach *in vivo*, removing any non-essential interactions via gene knockout and finely tuning the expression of the key components using promoter libraries^16,17^. This engineering approach – known as refactoring – makes a system easier to study by removing all non-essential natural regulation and feedback, thus enabling the system to be more efficiently tuned and directly measured. Effectively this generates cells streamlined for improved understanding of pathways and systems, while also making these cells more straightforward to utilize in downstream applications.

Here, we used genome engineering to construct a heavily-modified yeast suitable as an *in vivo* model for tuning GPCR signaling. By removing non-essential components, native transcriptional feedback regulation and all connections to the mating response we built a model strain retaining only the core signaling elements. In conjunction with a mathematical model, we used promoter libraries to vary the key components in this simplified, refactored pathway and revealed principles for tuning the sensitivity, leakiness and signal amplitude of the input-output dose-response curve. This new knowledge provides us with a rational approach for tuning signaling characteristics and, as we demonstrate, enables us to quickly reprogram yeast to sense and measure a variety of different inputs, either in single cell systems or community-based systems.

## Results

### A highly-engineered model strain for probing pathway variability

The pheromone response pathway is one of two native GPCR signaling pathways in *S. cerevisiae*^18^ and has long been the go-to choice for coupling heterologous GPCRs to yeast gene expression or for building systems for evolving GPCRs to desired targets^19,20^. Core to this pathway is an extensively-studied mitogen-activated protein kinase (MAPK) signaling cascade that functions with its own intrinsic feedback to maintain a robust input-output relationship in varying conditions^21^. As this MAPK cascade can be considered as a black-box processing unit in transduction through the pathway^11,22^, we chose to make this natural system the core from which we build and tune GPCR signaling pathways.

Keeping the six genes of the MAPK cascade fixed, we set out to generate a model strain for our work by first removing all other GPCR pathway-related genes from *S. cerevisiae* (**Figure 1**). This required making precise changes at 16 genomic loci in BY4741 yeast (see **Supplementary Table S1**), generating our model strain, yWS677, via eight rounds of CRISPR/Cas9-mediated editing (**Supplementary Figure S1**). Genomic changes were validated at each round by PCR and locus sequencing and all 16 changes in the final strain genome were verified by long-read nanopore sequencing (**Supplementary Figure S2**).

**Fig. 1.**
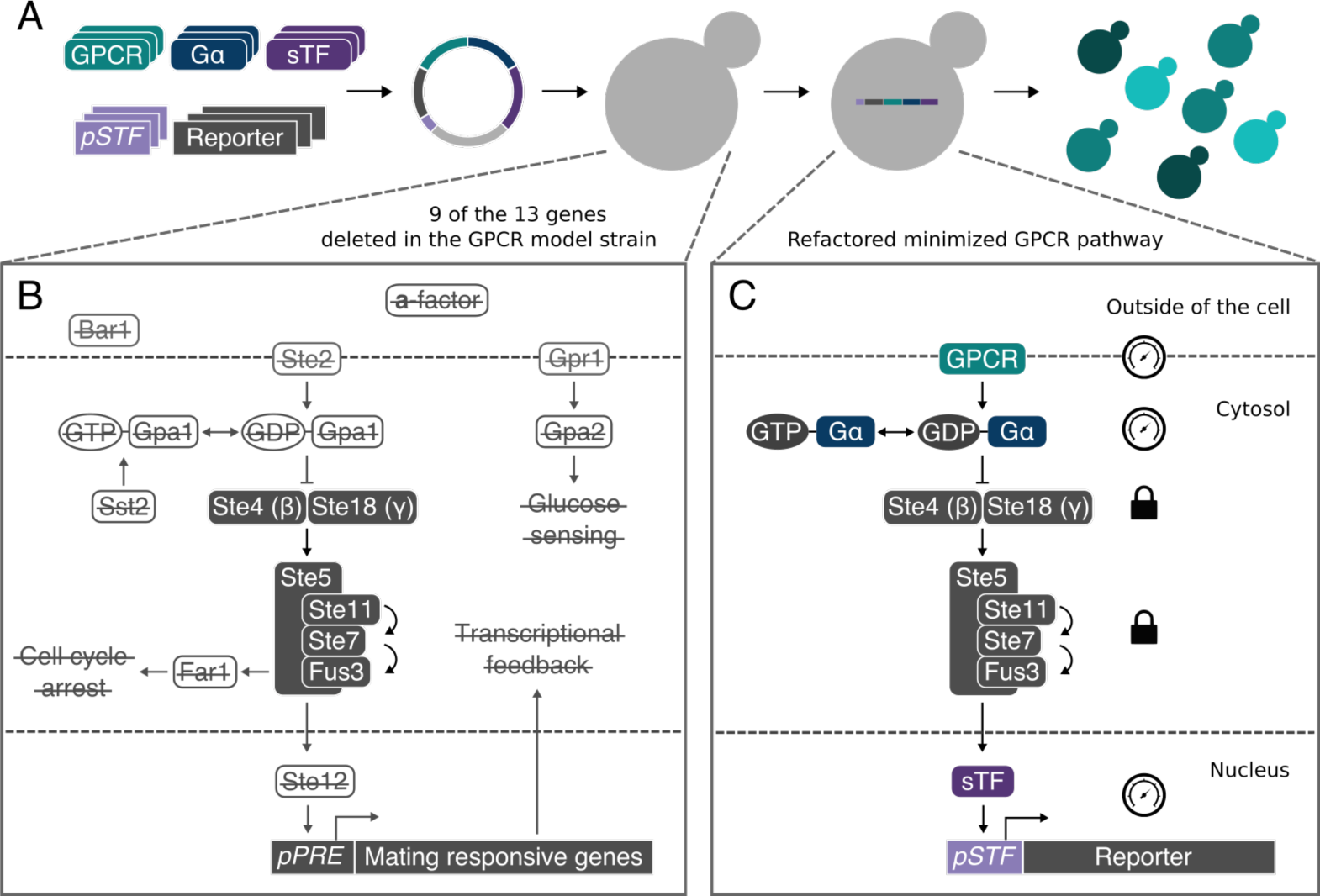
Model GPCR strain for probing pathway variability. (*A*) Pathway variants are generated by assembling parts from a library of signaling components into a single multigene cassette and then chromosomally integrating into the model strain, yWS677, to reconstitute a minimized GPCR signaling pathway. (*B*) 9 of the 13 genes deleted in the yeast mating and glucose-sensing pathways in the model strain, leaving only the Gβγ and core signaling element of the MAPK cascade. For a full list and description of all the changes see **Supplementary Table 1**. (*C*) A refactored signaling pathway, incorporating a heterologous receptor (GPCR) coupled to the pathway via a chimeric Gpa1-Gɑ subunit (Gα), and the output of the pathway redirected through a synthetic transcription factor (sTF) to an orthogonal promoter (*pSTF*).

During strain construction, we added addressable 24 bp targets in place of the open reading frames (ORFs) of all deleted genes. These allow for rapid and markerless (re)insertion of native or heterologous ORFs into these locations by CRISPR-aided multiplex integration (**Supplementary Figure S3**). For stable single-copy addition of further genes, three highly-characterized landing pads were also introduced that interface with rapid modular cloning system, Yeast MoClo Toolkit (YTK), which enables rapid multigene construction from high-characterized parts^23^. These changes were designed to facilitate rapid exploration of the effects of altering individual components of the pheromone response pathway.

To determine how the model strain performed with all non-essential components removed, the native receptor (*STE2*), Gα (*GPA1*), and pheromone responsive transcription factor (*STE12*) genes were reinserted at their natural loci to generate the “Quasi-WT” strain (**Figure 1C**). The α-factor dose-response of this strain was then compared to BY4741 yeast using the pheromone response *FUS1* promoter driving sfGFP expression^24,25^. As expected, a substantial shift in sensitivity and signal output was observed due to the minimization of the signaling pathway (**Supplementary Figure S4**).

### An *in vivo* model of the minimized pheromone response pathway

Previous work has shown that sensitivity of the yeast pheromone response pathway can be changed by altering the receptor number in the absence of the regulator of G protein signaling (RGS), Sst2^26^. Leakiness from constitutive receptor activity can also be reduced by overexpressing Gα^27,28^, as this acts as a negative regulator of signaling since Gβγ propagates the response to the MAPK cascade^9^. Therefore, to mathematically explore how sensitivity and leakiness could be varied by altering the expression levels of receptor and Gα, we built a single-cubic ternary complex model to capture the minimized pheromone response pathway upstream of the MAPK cascade^29^ (**Figure 2A+B**). We then systematically probed the response of the pathway in this model by individually altering the initial receptor and Gα concentrations, whilst keeping all other components fixed (**Figure 2C+E**). This demonstrated a clear relationship between the receptor number, sensitivity and maximum signal, as previously shown by Bush *et al*. (2016).

**Fig. 2.**
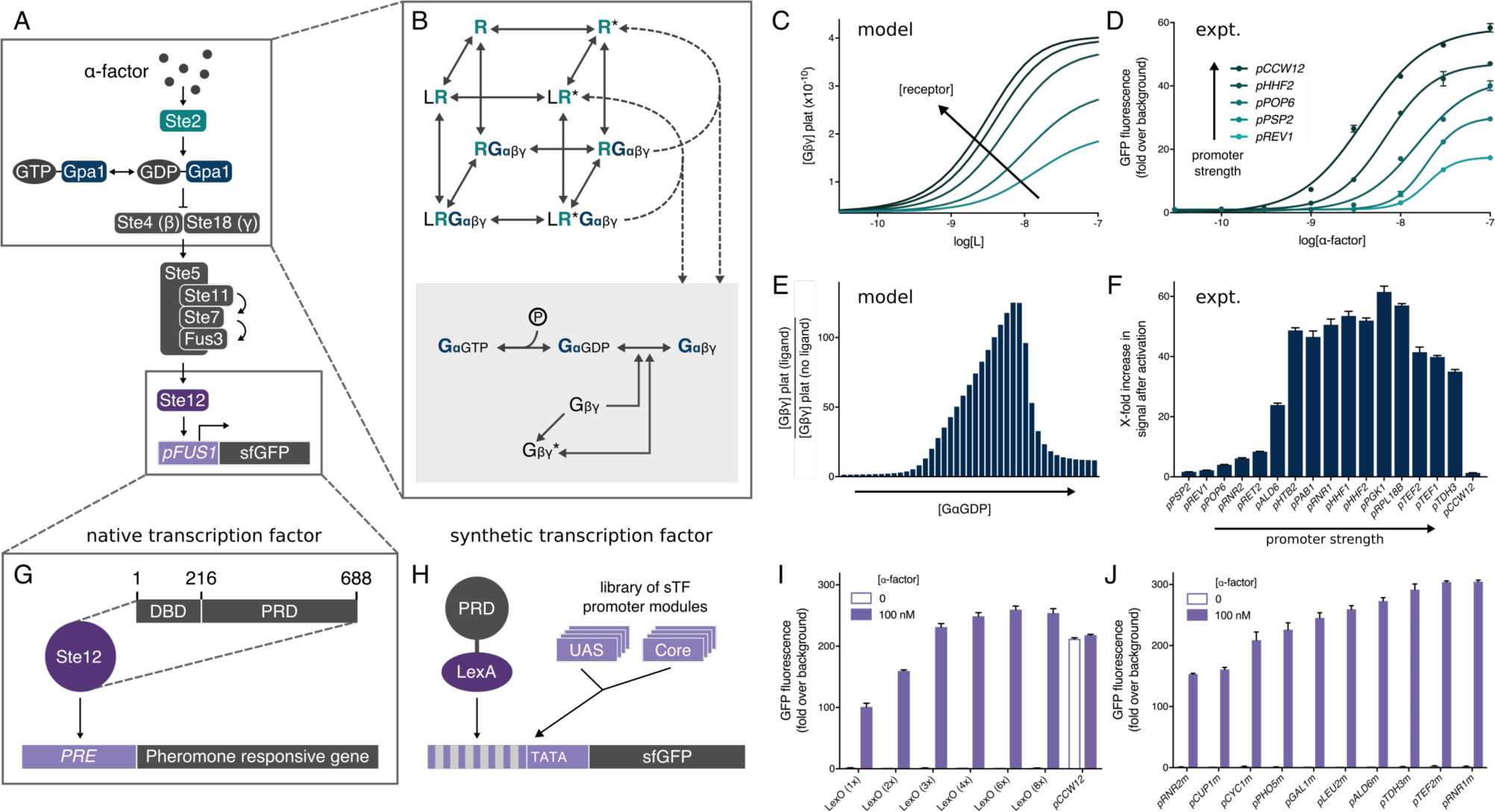
Varying the response of the minimized pathway by changing the promoter identity of key signaling components. (*A*) The minimized pheromone response pathway, composed of the native receptor, Gα, and pheromone-responsive transcription factor, under the control of constitutive promoters. (*B*) Single cubic ternary complex model of receptor/G protein signaling in the minimized pheromone response pathway. (*C*) Model of the pathway dose-response to α- factor over a range of initial receptor concentrations. (*D*) α-factor dose-response curves for 6 pathway designs, varying the expression of the receptor, Ste2, with a promoter library. (*E*) Model of maximum activation of the pheromone response pathway over a range of initial G_αGDP_ concentrations, demonstrating a predicted “sweet spot” of Gα concentration for which maximum fold-change in output is expected (See **Supplementary Figure 5** for extended model). (*F*) Maximum x-fold change in signal of a library of pathway designs, varying the expression of the Gα, Gpa1, with a promoter library. (*G*) Native pheromone-responsive transcription factor, Ste12, composed of a DNA binding domain (DBD; 1-215) and pheromone-responsive domain (PRD; 216-688), targeting a mating response gene, identified by a pheromone-response element (PRE). (*H*) Fusion of the full-length bacterial LexA transcriptional repressor with the pheromone-responsive domain controlling the expression of a modular promoter with an interchangeable UAS and core promoter, upstream of sfGFP. (*I*+*J*) Maximum α- factor-activated pathway expression mediated by the LexA-PRD transcription factor driving the expression from a synthetic promoter, using variants of the UAS and core promoter module, respectively. Data was acquired in the order defined in Figure 3. Results are means ± standard deviation from triplicate isolates.

The relationship between Gα levels and pathway response was more complex. At lower concentrations of Gα, constitutive activity of the pathway was observed due to increased free Gβγ. This, in combination with reduced receptor mediated signaling (due to lower receptor/Gαβγ concentrations) leads to a lower maximum-fold change in pathway activation. At higher Gα concentrations free Gβγ is rapidly sequestered, also leading to a decrease in the maximum fold change in pathway activity by acting as a “sponge” to signaling (**Supplementary Figure S5**). Thus, this model predicts a “sweet spot” of Gα expression between the two extremes, where all three members of the heterotrimeric G protein appear to be in balance, leading to a high-fold change in signal after activation.

Using the model strain and modular cloning system, we experimentally validated the findings of the mathematical model using a minimized pheromone response pathway with the receptor, Gα and transcription factor all rationally refactored (**Supplementary Figure S6**). In this approach, different constitutive promoters were selected to express the three components at native levels, and the *FUS1* promoter driving the expression of sfGFP was used as the response reporter. Once the minimized pathway had been validated (**Figure 3A, Design 1**), we then individually varied the expression of the receptor and Gα, while keeping all other components fixed. This gave results that qualitatively matched the model, demonstrating tuneability of the sensitivity through receptor number and revealing the predicted ‘sweet spot’ of Gα expression levels that gives the peak response (**Figure 2D+F**).

**Fig. 3.**
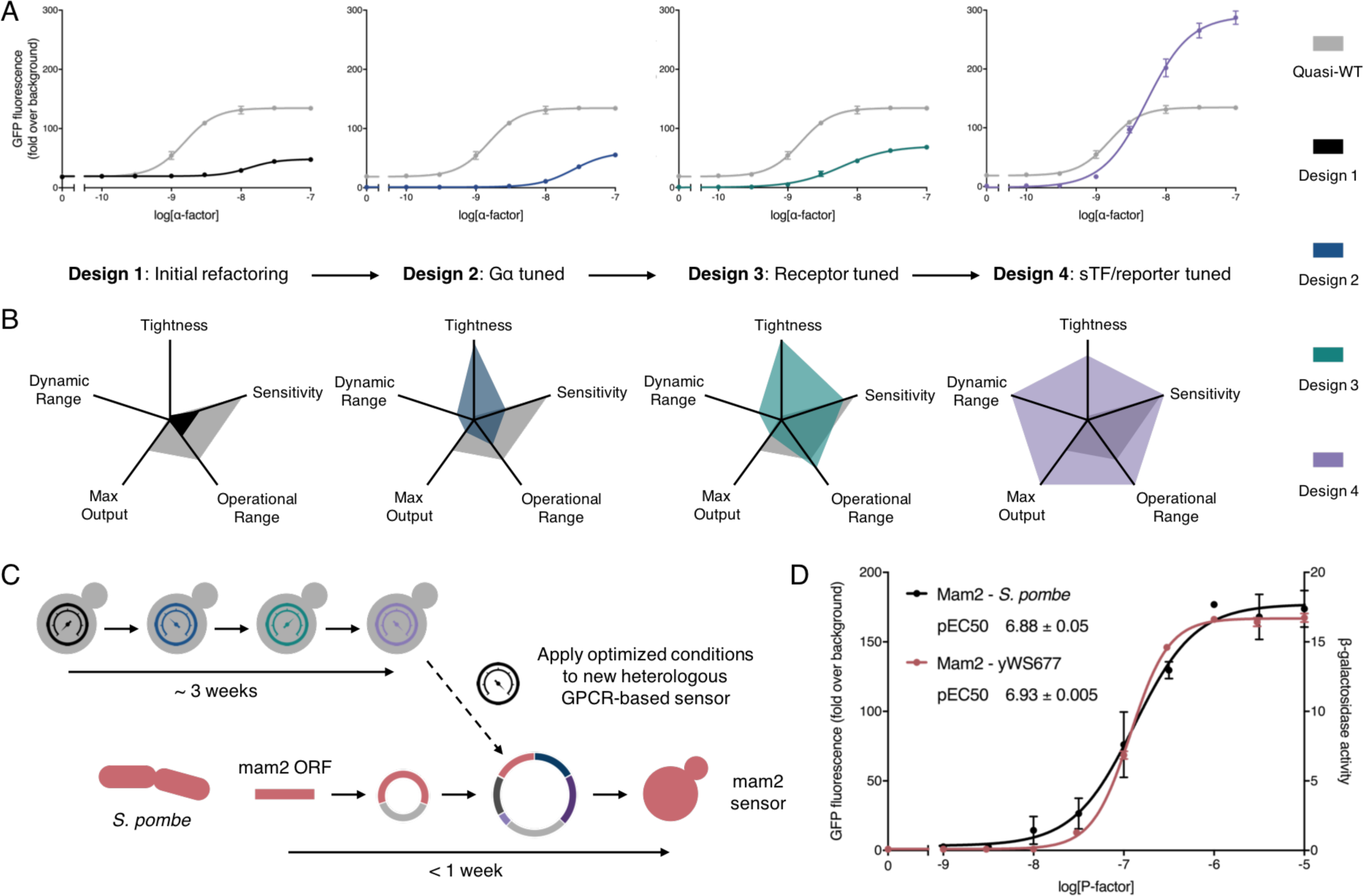
Tuning the minimized α-factor pheromone response pathway through iterative refactoring of the Gα, receptor, and sTF/reporter. (*A*) α-factor dose-response curves for the 4 sequential minimized pathway designs compared to the Quasi-WT response. (*B*) Dose-response characteristics for the 4 minimized pathway designs compared to Quasi-WT. For tightness, maximum output, and dynamic range, results were calculated from the measured minimum and maximum output. Sensitivity and operational range were determined from the fitted curve, defining sensitivity as the lowest 2-fold change over background, and operational range as the concentration span between the lowest and highest 2-fold change over background. All values were then normalized to the minimum measurable value and the maximum measured/calculated value within the dataset. For a table of the obtained values see **Supplementary Table 2**. (*C*) Domesticating the *S. pombe* Mam2 receptor in yWS677. The conditions identified during the 3-week optimization of the α-factor response, using the native Ste2 receptor, were directly applied to the design of the Mam2 sensor strain, which was built in less than a week. (*D*) P-factor dose-response curves of the Mam2 sensor (red) compared to the wild-type Mam2 response in its native *S. pombe* background (black) using previously characterized data from Croft *et al*. (2013)^35^. Slight differences in the shape of the curves are likely due to differences in the assay length and choice of reporter. Results are means ± standard deviation from triplicate isolates. Curves were fitted using GraphPad Prism variable slope (four parameter) nonlinear regression fit. *S. pombe* Mam2 dose-response data was kindly provided by Croft *et al*. (2013)^35^, and represents P-factor-dependent transcription of β-galactosidase using the sxa2 promoter, taking measurements 16 hours after stimulation.

Next, we sought to modulate the maximum pathway output by redirecting the pheromone-responsive transcription factor, Ste12, to synthetic promoters that can be tuned due to their modular nature^30^ (**Figure 2G+H**). Increasing the expression levels of the Ste12 transcription factor was not possible due to toxicity issues (**Supplementary Figure S7**), so we kept the expression of all transcription factor variants fixed at initial conditions. By fusing the pheromone-responsive domain of Ste12 (PRD; 216- 688) to the full-length LexA bacterial repressor protein^22,31^ we generated a synthetic transcription factor (LexA-PRD) able to target the pathway output to a library of modular synthetic promoters containing LexA operator sequences (LexO). These promoters were designed to direct different levels of expression by having different numbers of upstream transcription factor binding sites or different core promoter regions^30^. Both designs enabled us to vary the maximum output of the response over a three-fold range, without compromising the tightness of the OFF state when the pathway is not activated (**Figure 2I+J**). Externally tuning the maximum pathway output was also possible using ligand-inducible DNA binding domains (DBDs) fused to the PRD^32,33^ (**Supplementary Figure S8**).

Finally, we validated the decoupling of the pheromone response pathway and the mating response in our engineered strain and confirmed the orthogonality between synthetic transcription factors and their target promoter pairs (**Supplementary Figure S9**).

### Rational optimization of α-factor and P-factor sensing through refactoring

Together our mathematical and *in vivo* models reveal that GPCR dose-response variability can be achieved by altering the promoter identify for just three components (receptor, Gα, and reporter), offering a simple approach to rationally tune the sensitivity, leakiness, and signal output of a GPCR pathway.

To demonstrate this in practice, we next optimized the α-factor response of our minimized response pathway through iterative refactoring of the key components (**Figure 3A+B**). Our starting strain, with constitutive expression of the three components set at native levels (**Design 1**), performed poorly compared to the Quasi-WT strain due to the removal of the native regulated promoters which incorporate feedback control (**Supplementary Figure S11**). However, increasing Gα levels reduced the leakiness of the response to effectively zero (**Design 2**). This was done by using the *PGK1* promoter, which gave the desired raised levels but crucially avoided overexpressing Gα in a way that would reduce pathway sensitivity when coupled to different receptors (**Supplementary Figure S12**). The dose-response sensitivity could then also be boosted by increasing receptor expression via the strongest promoter available (*pCCW12*). This version (**Design 3**) now approached the sensitivity of the Quasi-WT strain for the α-factor inducer. Finally, the output strength of the response could be optimized by linking the pathway to activate the best performing synthetic promoter, *LexO(6x)- pLEU2m*, via the LexA-PRD synthetic transcription factor (sTF). The resulting pathway (**Design 4**) was highly optimized in comparison to the Quasi-WT strain with improved operational range, tightness, dynamic range and maximum output. Notably, this is achieved without the need for feedback regulation of signaling components and also offers a pathway decoupled from the >100 genes usually upregulated in the mating response^34^.

Understanding how the pathway dose-response can be shifted in this manner advances our basic knowledge of how component level changes effect signal transduction. Alongside this, it also offers direct applications for synthetic biology, where reprogramming cells to receive specific signals and respond in a desired manner is a core goal^36^. Indeed, GPCRs represent the ideal sensory module for eukaryotic synthetic biology as they are responsive to a plethora of ligands and stimuli, often operate with high specificity^37^, and naturally have modularity written into their signaling architecture^38,39^.

With this in mind, we next turned to utilizing our model strain and toolkit for rationally engineering yeast cells as biosensors that sense diverse inputs via heterologous GPCRs (**Supplementary Figure S10**). As an initial demonstration we took the Mam2 receptor from *Schizosaccharomyces pombe*, which detects a 23 amino acid peptide called P-factor^40^. Following the optimized tuning levels determined above for Design 4, we generated a P-factor-sensing strain in less than a week. We then compared the response of this sensor strain to the response observed in its native context, as reported by Croft *et al*. (2013)^35^ (**Figure 3D**). The Mam2 sensor strain behaved almost exactly as in *S. pombe*, achieving an almost identical potency (pEC50) to P-factor. Furthermore, the Mam2 sensor strain displayed no detectable leakiness and exhibited a 180-fold change in signal after activation, suggesting the optimization we had performed would also be suitable for other GPCRs.

### Mixed populations for extending and narrowing the operational range of GPCR-sensing

Efforts to create sensor strains over the last two decades have coupled heterologous GPCRs to the yeast pheromone response pathway with varying success. Now with the approach described here the sensitivity, leakiness and response output of cells sensing via heterologous receptors can be rationally tuned. However, one further important characteristic of a sensor - its operational range (the Hill slope of its dose-response curve) - is more difficult to adjust, as it is determined largely by the ligand-binding properties of the receptor. Some receptors will confer a narrow range switch-like behavior, only requiring a small increase in signal to trigger maximum output (*i.e.* a digital response), whereas others will give a wide operational range where there is a proportional relationship between signal and output (*i.e.* a linear response)^41^. For sensor applications, a linear response is typically required, whereas the digital response is more desirable for gene circuit applications.

With this in mind, we next set-out to solve how the operational range can also be tuned using an engineering approach, so that the Hill slope for a digital-like sensor can be reduced to expand the range, while the Hill slope can be increased on a linear-like sensor to narrow the operational range. For this we first built two new sensor strains, both sensing medically-relevant metabolites by having human GPCRs coupled to our refactored yeast pathway. The chosen receptors were the adenosine-responsive A2BR receptor, previously shown to give a digital-like response in yeast^42^, and the melatonin-responsive MTNR1A receptor, previously shown to give a linear-like response in yeast^43^ (**Supplementary Figure S13**).

While previous efforts have tuned the yeast pheromone response Hill slope by overlaying synthetic feedback loops into the MAPK cascade^44–46^, feedback loops on the components tuned in our approach proved ineffective in changing the dose-response curves (**Supplementary Figure S14**), likely because our orthogonal sTFs do not have the autoregulatory feedback that Ste12 has via its native promoter^47^. Without this avenue, we instead choose a different tactic; tuning the Hill slope by creating engineered communities of cells that change the average response at the population level.

Firstly, to create a population that linearizes the steep response of our adenosine-sensing cells, we took inspiration from a strategy employed by previous artificial biosensor systems, where receptors with different sensitivities are combined and their average response determines the output^48^. We used rational tuning of receptor and Gα levels to create two new strains with increased and decreased sensitivity to adenosine, and then tuned these at the output promoter so that their maximum outputs match the original strain (**Supplementary Figure S15**). We then co-cultured the three-sensing strains in a 1:1:1 ratio to create a consortium whose average response integrates the signal from all cells to give an extended operational range (**Figure 4A**). This almost halved the Hill slope of the response whilst maintaining a similar potency, yielding an operational range 50-fold greater than the initial response (**Figure 4B-D**).

**Fig. 4.**
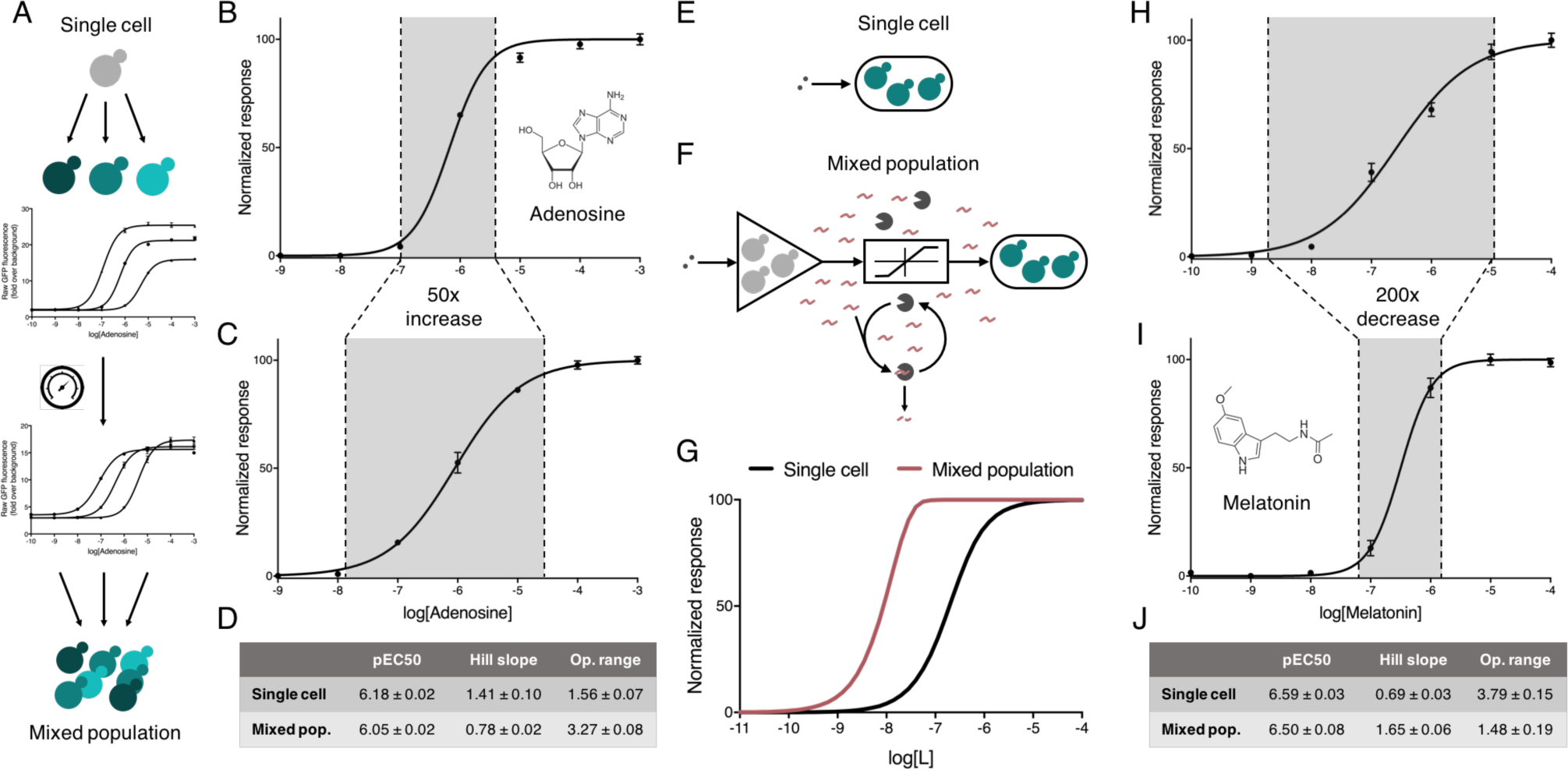
Mixed populations of cells for extending or narrowing the operational range of heterologous GPCR sensors. (*A-D*) Engineered populations for extending the operational range of adenosine sensing. (*A*) Engineered cells combined to produce a system with an extended operational range. First, a range of cells are produced with different sensitivities to a particular ligand by expressing the receptor to varying degrees. Next, the ligand responses are balanced to produce similar maximum outputs. Finally, the cells are combined in equal parts to create a mixed population of cells with an extended operational range. (*B*) The limited dose-response of the human A2BR receptor to adenosine in a single cell line, operational over 1.6 orders of magnitude. (*C*) The extended dose-response of a population of engineered cells to adenosine, operational over 3.3 orders of magnitude. (*D*) Potency (pEC_50_) and Hill slope values for the single and mixed population A2BR sensor strains. (*E-J*) Communicating cells for narrowing the operational range melatonin sensing. (*E*) An MTNR1A sensor strain. (*F*) A mixed population of amplifier and reporter cells designed to create a digital response from an otherwise linear sensor. In response to ligand, amplifier cells release α-factor which is detected by reporter cells constitutively secreting the α-factor degrading protease, Bar1. The presence of Bar1 degrades low levels of α-factor preventing reporter strain activation until levels of α-factor are high enough to saturate the capacity of Bar1-mediated degradation. (*G*) Model of the amplifier-reporter system with Bar1-mediated threshold response. (*H*) The broad dose-response of the human MTNR1A receptor to melatonin in a single cell line, operational over 3.8 orders of magnitude. (*I*) Digitization of melatonin sensing, reducing the operational range of the response to 1.5 orders of magnitude, 200-fold lower than a single population. (*J*) Potency (pEC_50_) and Hill slope values for the single cell and mixed population sensor systems. Here, the operational range is defined as the difference between 5% and 95% of the activated response. Results are means ± standard deviation from triplicate isolates.

Narrowing the operational range of the melatonin-responsive MTNR1A sensor strains required more complex engineering as the Hill slope of a response can only be increased via mechanisms such as cooperativity^41^, sequesteration^49^ or positive feedback^50^. As before, we utilized a community-based approach, but here using cell-to-cell communication to enable feedback at the population level^51,52^. A two-cell system was designed where the first cell acts as an amplifier, sensing via MTNR1A and responding by secreting α-factor from the reintroduced gene. The second cell senses α-factor and responds with reporter gene (sfGFP) expression, and also secretes constitutive levels of the α-factor degrading protease, Bar1, to create a threshold for activation (**Figure 4F**). Computational modelling of this system (**Figure 4G**) identified that fine-tuned expression of Bar1 is key to producing a steep Hill slope while maintaining low leakiness and high dynamic range (**Supplementary Figure S16**).

Following the creation of the α-factor-detecting reporter cells and the tuning of Bar1 levels secreted by these, we engineered the digital response in two steps using our toolkit (**Supplementary Figure S17**). Amplifier cells were first created by linking the output of MTNR1A sensor strain to the production of peptide pheromone, α-factor. The sensitivity of these cells was then adjusted so that the potency of the response in the final two-cell system matched that of the MTNR1A response in the single cell system by adjusting the MTNR1A receptor expression in the amplifier strain. When co-cultured in a 1:1 ratio, the final tuned two-cell system maintained the same potency but now provided a dose-response curve with a 2.3x increase in Hill slope and more than 200-fold decrease in the operational range (**Figure 4H-J**).

## Discussion

In this study, we used genome engineering and synthetic biology tools to refactor a minimal GPCR signaling pathway so that it could be rationally tuned both *in silico* and *in vivo*. This revealed simple mechanisms for tuning the sensitivity, leakiness and signal amplitude of the pathway dose-response curve, which enabled us to engineer yeast strains for desired performance as biosensors for peptide inducers, and for primary and secondary metabolites.

The refactoring approach made it trivial to treat our strain as an *in vivo* model of a pathway, as performance was simple to measure from fluorescent reporter output, and the expression of key components could be individually varied while all other components remain fixed. This makes it highly analogous to an *in silico* model system, and led us to new insights into the importance of protein stoichiometries in GPCR pathways. Notably we showed that changing the promoter strengths for the receptor, Gα, and reporter all significantly altered the dose-response in predictable ways that held for all four tested receptors. This indicates the generic applicability of the tuning principles uncovered here, and also demonstrates that each GPCR-mediated response is not merely defined by the receptor’s intrinsic properties (i.e. ligand affinity); instead it is a function of the properties of all components in the signaling pathway and particularly their relative levels.

This fact has important consequences. It explains how signal transduction behavior could be significantly altered by a change in component levels, whether due to a change in environmental conditions or due to altered expression and protein turnover in different tissues. Indeed, via this mechanism cells can have different sensitivities and activation thresholds for the same agonist whilst expressing identical receptors. Importantly, this fact also underlines why non-coding genetic variation, such as mutation in promoter regions, has to also be considered alongside protein polymorphisms when assessing how genetic variation links to health and to the efficacy of treatments^53^. Already, receptor variation in humans is recognized as a major cause of GPCR-targeting drugs being ineffective in many individuals^54^, and it is possible that non-coding mutations that alter pathway stoichiometries can further explain many cases. The importance of stoichiometry may also explain failures from past efforts in pharmacology research to characterise cloned GPCRs by expressing these at high copy from strong promoters in yeast or mammalian cell lines. Overexpression of a receptor without considering the need for tuning the G protein will typically lead to a leaky system with high basal activation without any agonist present^55^.

We anticipate that the tuning principles uncovered here in yeast will also be relevant for GPCR signaling in all eukaryotes, however, it is worth re-stressing the large diversity in the type and structure of downstream signaling pathways paired with GPCRs in different organisms and cell types. The next steps for our approach will therefore be to use equivalent tools to refactor a canonical mammalian GPCR pathway so that its components can be tuned and assessed in isolation to the point where the dose-response to the agonist can be set as desired. This would also accelerate applications in pharmacology and healthcare that utilize GPCRs, such as in cell-based theranostics where cells are engineered to detect and act upon receiving defined cues within the human body^37^. Moreover, the mammalian system will require tuning on not one G Protein but the many endogenous Gα that exist, all of which could interact to influence the activity seen via the others^8^. It is also possible that this complicated orchestra plays a role in ligand bias where it is probable that the output seen is not only a function of the receptor conformation but the type and number of G protein available in a specific cell type^56^.

As we have demonstrated here, our model strain also offers immediate applications for engineering yeast as biosensors, whether as single strains or as part of engineered consortia. Already efforts in synthetic biology have used engineered yeast biosensors as medical diagnostics^57^, for pathogen detection^58^, and as a tool for accelerating metabolic engineering^22,59^. In all these applications, it is desirable for the user to have control over the response to input and the magnitude of gene expression it triggers. Full ability to tune biosensors, as shown here, allows engineering for desired detection windows, and could be used in further work to define thresholds for activation (e.g. for directed evolution^20,57^) or for matching the input/output levels when cells are engineered to detect and act, or to communicate in connected systems^60^.

A current limitation of using yeast for biosensors is that most medically-relevant GPCRs do not port directly into *S. cerevisiae* without requiring optimization of expression, membrane translocation and pathway coupling. Solutions to this have previously been explored with limited success, such as the co-expression of mammalian accessory proteins^61^ and the humanization of the yeast membrane^62^. Nevertheless, with the four receptors we have shown to couple successfully in this system, the principles of signal tuning are maintained across all of them. In doing so we have established a fundamental hypothesis for one of the biology’s most important cell communication systems.

Our model strain now offers a starting point to develop work towards this in a more systematic, plug-and-play manner. By starting with a low-complexity, insulated pathway with direct output, a bottom-up approach can be used to predictably increase capabilities. Indeed, a next step would be to introduce control over the signaling dynamics and feedback by also refactoring the well-characterized MAPK pathway core left untouched here; in tandem also incorporating established models of this into the accompanying *in silico* model. The overall strategy, of simplifying and refactoring a natural pathway to first understand it and then predictably expand it, should also be applicable to other systems in and beyond signal transduction. With the accelerating capabilities of genome engineering and synthetic biology in all organisms, it is likely that we will see the creation of equivalent *in vivo* model strains to rationally explore and exploit the key features and parameters of other important biological systems.

## Acknowledgements

The authors wish to thank Dr Benjamin Blount and Dr Charlie Gilbert for advice and input during this project and acknowledge the UK Biotechnology and Biological Sciences Research Council (BBSRC) for the funding of this work as a CASE PhD award in collaboration with AstraZeneca.

## Materials and Methods

### Strains and Growth Media

The *S. cerevisiae* parental strain used for all experiments was yWS677 (MATα his3Δ1 leu2Δ0 met15Δ0 ura3Δ0 sst2Δ0 far1Δ0 bar1Δ0 ste2Δ0 ste12Δ0 gpa1Δ0 ste3Δ0 mf(alpha)1Δ0 mf(alpha)2Δ0 mfa1Δ0 mfa2Δ0 gpr1Δ0 gpa2Δ0), a strain engineered within this study from BY4741 (MATα his3Δ1 leu2Δ0 met15Δ0 ura3Δ0). To generate the Quasi-WT strain and GFP-ORF substitution strains, yWS677 was reverse edited to either restore the STE2, GPA1, and STE12 to wild-type in one cell line or substitute with the sfGFP ORF in separate strains.

All experiments were performed in synthetic complete media with 2% (w/v) glucose (VWR), 0.67% (w/v) Yeast Nitrogen Base without amino acids (Sigma), 0.14% (w/v) Yeast Synthetic Drop-out Medium Supplements without histidine, leucine, tryptophan, and uracil (Sigma), 20 mg/L histidine (Sigma), 100 mg/L leucine (Sigma), 20 mg/L tryptophan (Sigma), 20 mg/L uracil (Sigma). YPD was used for culturing cells in preparation for transformation: 1% (w/v) Bacto Yeast Extract (Merck), 2% (w/v) Bacto Peptone (Merck), 2% glucose (VWR).

NEB^®^ Turbo Competent *E. coli* was used for all cloning experiments. Transformed cells were selected on Lysogeny Broth (LB) with the appropriate antibiotics (ampicillin 100 μg/mL, chloramphenicol 34 μg/mL, or kanamycin 50 μg/mL).

### Yeast Transformations

Yeast colonies were grown to saturation overnight in YPD, then diluted 1:100 in 15 mL of fresh YPD in a 50 mL conical tube and grown for 4-6 h to OD600 0.6-0.8. Cells were pelleted and washed once with 10 mL 0.1 M lithium acetate (LiOAc) (Sigma). Cells were then resuspended in 0.1 M LiOAc to a total volume of 100 μL/transformation. 100 μL of cell suspension was then distributed into 1.5 mL reaction tubes and pelleted. Cells were resuspended in 64 μL of DNA/salmon sperm DNA mixture (10 μL of boiled salmon sperm DNA (Invitrogen) + DNA + ddH_2_O), and then mixed with 294 μL of PEG/LiOAc mixture (260 μL 50% (w/v) PEG-3350 (Sigma) + 36 μL 1 M LiOAc). The yeast transformation mixture was then heat-shocked at 42°C for 40 mins, pelleted, resuspended in 200 μL 5 mM CaCl_2_ plated onto the appropriate solid agar media.

### Iterative Markerless Editing of Yeast Genome

All genomic edits were performed using CRISPR/Cas9 mediated genome editing and validated by colony PCR followed by sanger sequencing of the amplified genomic region (for a list of primers used in this study, see **Supplementary Table S6**). To rapidly iterate between successive edits when generating the model strain yWS677, a marker cycling protocol was used, where two deletions or changes were performed per round of editing. Plasmid curing was skipped, and the marker for selecting the CRISPR/Cas9 plasmid was cycled between 3 markers (*URA3*, *LEU2*, and *HIS3*) for each iteration. The final edit was performed with CRISPR/Cas9 plasmid containing the *URA3* selection marker, which was counter selected using 5-FoA to cure the yeast of CRISPR machinery. The absence of all CRISPR/Cas9 plasmids was validated by colony PCR and replica plating on selectable media.

Protospacer sequences for sgRNAs targeting the genome and generating new and unique landing pads were designed using Benchling (see **Supplementary Table S1** and **Supplementary Table S5** for a list of LPs and sgRNAs used in this study). All sgRNAs were cloned into pWS082 sgRNA entry vector. For more information on donor DNA design see **Supplementary Figure S1**. For more information on the toolkit and protocol used to iterate successive edits see https://benchling.com/pub/ellis-crispr-tools.

### Nanopore Sequencing of the yWS677 Genome

DNA was isolated from yWS677 for Nanopore sequencing using the 100/G Genomic-Tip kit (QIAGEN), sheared to 20 kb using a g-TUBE (Covaris) and prepared for sequencing using a Ligation Sequencing Kit 1D^2^ R9.5 (Oxford Nanopore Technologies). The genomic DNA was then run on an R9.5 flow cell using a MinION Mk 1B (Oxford Nanopore Technologies). A standard 48h sequencing run was performed using the MinKnow 1.5.5 software using local basecalling. Reads were exported directly to fastq using MinKNOW. Canu (v1.5) was used to correct raw reads (www.canu.readthedocs.io) and smartdenovo (www.github.com/ruanjue/smartdenovo) was used to de novo assemble the reads into contiguous sequences (contigs) using default flags. Resulting contigs were compared to a WT reference genome (s288c, SGD) using lastdb/lastal (www.last.cbrc.jp) and viewed on integrative genome viewer (IGV) (www.software.broadinstitute.org) to inspect genomic changes.

### Plasmid Construction

All plasmids within this study were created using the Yeast MoClo Toolkit (YTK)^23^. Additional sequences not included within the YTK toolkit that were used within this study can be found in the **Part sequences** in the Supplementary Information. For more information on using the Yeast MoClo Toolkit see Lee *et al*., (2015). For more information on the plasmid formatting and a list of all plasmid constructs used in this study, see **Supplementary Figure S10** and **Plasmids used in this study** in the Supplementary Information. Part sequences were either mutated or designed and synthesized to remove or avoid all instances of the BsmBI, BsaI, BbsI, and NotI recognition sequences.

### Golden Gate Assembly Protocol

All plasmids for Golden Gate reactions were set to equimolar concentrations of 50 fmol/μL (50 nM) prior to experiments. Golden Gate reactions were prepared as follows: 0.1 μL of entry vector (backbone), 0.5 μL of each DNA plasmid, 1 μL T4 DNA ligase buffer (Promega), 0.5 μL T7 DNA Ligase (NEB), 0.5 μL restriction enzyme (BsaI or BsmBI) (NEB), and water to bring the final volume to 10 μL. Reaction mixtures were then incubated in a thermocycler using the following program: (42 °C for 2 min, 16 °C for 5 min) x 25 cycles, followed by a final digestion step of 60 °C for 10 min, and then heat inactivation at 80 °C for 10 min.

### High-Efficiency Multiplexed Integrations

The yWS677 model strain was prepared for multiplex integration of marker cassettes by integrating landing pads (LPs) at the *URA3*, *LEU2*, and *HO* loci, conforming to the YTK integration plasmid format. Using CRISPR/Cas9 targeting the unique protospacer sequences at the LPs of URA3, LEU2, and HO, the efficiency of integrating multiple selectable cassettes was greatly increased (**Supplementary Figure 3**).

Single, double, and triple integration of marker plasmids were performed with 50, 100, and 200 ng of plasmid, respectively, with 100 ng of Cas9 and 200 ng of each gRNA expression cassette. All plasmids were first linearized by digestion before transformation using NotI-HF (NEB). Successful plasmid integration was selected for using synthetic drop-out media missing the appropriate supplements. Cas9 and gRNA expression was transient and quickly lost due to lack of selection or homology to genome. Initially validated by colony PCR and then assumed thereafter.

### RT-qPCR

All quantitative PCR (qPCR) qPCR reactions were performed in an MasterCycler ep RealPlex 4 (Eppendorf) using SYBR FAST Universal qPCR Master Mix (Kapa Biosystems) according to the manufacturer’s instructions. For RNA purification, RNA was isolated from yeast culture grown to an OD600 of 1 ± 0.1 using a YeaStar RNA Kit (Zymo Research) according to the manufacturer’s instructions. RNA was quantified by nanodrop spectrophotometer (Thermo Fisher) and cDNA was generated from each RNA prep using a High Capacity cDNA Reverse Transcription Kit (Applied Biosystems). Each qPCR reaction contained 20 ng of cDNA. qPCR results were normalized to the housekeeping gene *HTB2*. All qPCR primers were designed manually using Benchling.

### Ligand Sensing Protocol

All sensor strains were picked into 500 μL of synthetic complete media and grown in 2.2 mL 96 deep-well plates at 30 °C in an Infors HT Multitron, shaking at 700 rpm overnight. The next day, saturated strains were then diluted 1:100 into fresh media. After 2 h of incubation the strains were induced with their respective ligands and incubated for a further 4 h. All ligands were dissolved in DMSO, and the final concentration in all cultures was 1%. For strains using the Z_3_E-PRD and TetR-PRD transcription factor, aTc and β-estradiol was added during the back dilution at time 0 h. To perform flow cytometry and plate reader measurements, 200 μL from each well was directly transferred to a 96-well clear, flat-bottom microplate (Corning).

For monoclonal cell experiments, cell fluorescence was measured by an Attune NxT Flow Cytometer (Thermo Scientific) with the following settings for measuring sfGFP: FSC 300 V, SSC 350 V, BL1 500 V. Fluorescence data was collected from 10,000 cells for each experiment and analyzed using FlowJo software. For polyclonal cell experiments, cell fluorescence was measured by a Synergy HT Microplate Reader (BioTek) with the following settings for measuring sfGFP: excitation 485/20, emission 528/20, gain 80. Unnormalized, raw fluorescence readings from each well were used for data analysis.

### Dose-Response Fitting

All presented dose-response fittings were generated in GraphPad Prism 7, which was used to determine the pEC50 and Hill slope. To determine all remaining properties of the dose-response curve, curve fitting was performed using Python (SciPy and Matplotlib) using the 4PL model, where x is the concentration, A is the minimum asymptote, B is the steepness, C is the inflection point and D is the maximum asymptote:

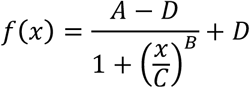

### Computational Modelling

Chemical kinetic models based upon reactions schemes shown in Supplemental Information were generated as a systems of ordinary differential equations (ODEs). The system of ODEs was solved using MATLAB R2017a offered by MathWorks. Data fitting, parameter estimation and model analysis were performed in COPASI. The full ODE equations, kinetic rates and simulations for each model are described in Supplementary Information.

